# Designing a Novel 3D Scaffold for Multiepitope Vaccine Development: Engineering Ag85a Protein for Enhanced Stability and Antigenicity

**DOI:** 10.1101/2024.03.20.585912

**Authors:** Torsha Mondal, Shakilur Rahman, Amit Kumar Das, Ditipriya Hazra, Amlan Roychowdhury

**Affiliations:** Department of Bioscience, JIS University, Kolkata, India; Department of Biotechnology, Indian Institute of Technology Kharagpur, India; Department of Biotechnology, St. Xavier’s College Kolkata, India; Center for Healthcare Science and Technology, Indian Institute of Engineering Science and Technology, Shibpur, India

**Keywords:** Protein engineering, Molecular dynamics simulation, reverse vaccinology, computational immunology, Tuberculosis

## Abstract

Designing multi epitope vaccine (MEV) by reverse vaccinology has become immensely important in the area of vaccine research due to the emergence of new pathogens as well as rise of multi drug resistant old evils like tuberculosis. Administering a vaccine may have the best possibility to save mankind from these unforeseeable events. The strategy of designing a MEV in-silico lies in a few basic steps, including procuring the amino acid sequence of the B cell and T cell epitopes from literature search, bioinformatics approach to construct a potent immunogen capable of eliciting both humoral and cell mediated response. But the challenge lies in the construction of a stable protein with a compact tertiary structure. Merely joining the epitopes one after another, may not be sufficient to achieve this. In this study, a methodology has been detailed to tackle this great challenge using a simple approach of protein engineering. A scaffold based MEV has been designed for the very first time against Mtb by converting a vaccine candidate protein, Ag85A into a scaffold by truncating its non-immunogenic regions so the gaps could be filled by the highly immunogenic epitopes. The stability of the MEV was estimated by molecular dynamics simulation.

## INTRODUCTION

Prediction of vaccine target, based on the genome sequence of the organism, also known as Reverse vaccinology, is able to predict robust vaccine candidates, that does not rely on growing the pathogen *in-vitro* [1]. Applied for the first time, for identification of vaccine candidates of *Neisseria meningitidis*, this technique has gained popularity in recent times for predicting vaccine candidates more efficiently than the existing techniques [2, 3]. Reverse vaccinology relies on computational tools to predict B-cell and T-cell epitopes that are highly immunogenic, from the genome sequence of the organisms. These epitopes are then linked to produce a multi-epitope vaccine that would be highly immunogenic since it elicits both humoral and cell mediated immune response, addition of an adjuvant to this multi-epitope vaccine (MEV) by a linker, provides long lasting immune response [4].

In modern world vaccines have become an indispensable first line of defence against life threatening infectious diseases. One of such deadly disease that is threatening mankind for centuries is tuberculosis (TB), caused by *Mycobacterium tuberculosis* (Mtb). In spite of the availability of a very well-known live-attenuated type vaccine (BCG), antibiotics and intensive care facilities; TB claimed the life of 1.6 million people worldwide in the year 2021 alone (according to WHO’s report) [5]. Considering this grave situation an attempt has been made using protein engineering, bioinformatics, computational immunology and molecular dynamics simulation to develop a unique method for the scaffold based MEV construction. Where Mtb has been considered as the model organism.

In this study we have used an approach to construct a scaffold based MEV against *Mycobacterium tuberculosis*, that is not only highly immunogenic but also more stable than conventional MEVs.

Global efforts to control tuberculosis are still seriously threatened by the emergence of multidrug resistance and HIV/TB co-infection. Even with incredible advances in medicine and science, tuberculosis treatment remains a challenge, due to evolution of drug resistant strains, late detection, dormant infection, and poor public health. The current treatment of tuberculosis includes first line of oral drugs, isoniazid, rifampicin, ethambutol and pyrazinamide, second-line injectable drugs, Streptomycin, Kanamycin, Amikacin, Capreomycin, along with Fluoroquinolones [6].

The most effective way to control tuberculosis is through vaccination. Till date only one vaccine exists against TB, Bacille Calmette Guerin (BCG), which is an attenuated vaccine, obtained from subculture of *Mycobacterium bovis* [7]. BCG provides protection from childhood tuberculosis and disseminated TB, like miliary TB, however, with time the efficiency of BCG fades, and provides very little protection against adult pulmonary TB, which results in spreading of TB among susceptible individuals [8]. Naturally, persistence of TB infection, despite BCG vaccination calls for more effective vaccines. Development of a new TB vaccine for adolescents and adults, can significantly lower the spread of TB, globally [9].

One of the major vaccine candidates is Antigen 85A subunit (Ag85A) of *Mycobacterium tuberculosis*. Ag85A is essential for the survival of the bacteria in host cell macrophages [10]. Studies have indicated Ag85A elicits a strong immune response [11, 12]. Currently, two adenovirus based vaccines, Ad5 Ag85A and ChAdOx1 85A-MVA85A, using Ag85A as the immunogen, are under trial [13–15].

From extensive literature survey, seven secretory and cell surface proteins of *Mycobacterium tuberculosis* were selected, RV3810, RV0101, RV1161, RV3870, RV3646c, RV1918c. RV3810 is a cell surface protein, essential for survival of the bacteria in host cell [16, 17]. RV0101 is critical for lipid metabolism, and RV1161 is required for nitrate metabolism and respiration [18, 19]. RV3870 plays a key role in pathogenesis, and RV3646c, RV1918c are upregulated in virulent strains [20]. These selected proteins were screened using computational immunology methods, to predict B-cell, Helper T-Lymphocyte (HTL) and Cytotoxic T-Lymphocyte (CTL) epitopes and these epitopes were again tested for antigenicity and toxicity, as well as allergenicity. The HTL and CTL epitopes were additionally screened on the basis of MHCI and MHCII binding. The epitopes selected from this study were used to construct two different kind of multiepitope vaccines, MEV1(multi epitope vaccine 1), which is a conventional MEV where epitopes are joined by linkers along with an adjuvant, and MEV6.2 (multi epitope vaccine 6, version 2) where the epitopes replaced secondary structures of a scaffold designed from Ag85A crystal structure.

In this study we have used a novel protein engineering approach where Ag85A has been used as a scaffold. The flexible and non-immunogenic regions of Ag85A have been replaced with the selected epitopes to obtain MEV6.2 (multi epitope vaccine 6.2). The strategy is summarised graphically in Figure 1. The stability was assessed by, molecular dynamics simulation (MD simulation) and the efficacy as immunogen by immune simulation.

**Figure 1:**
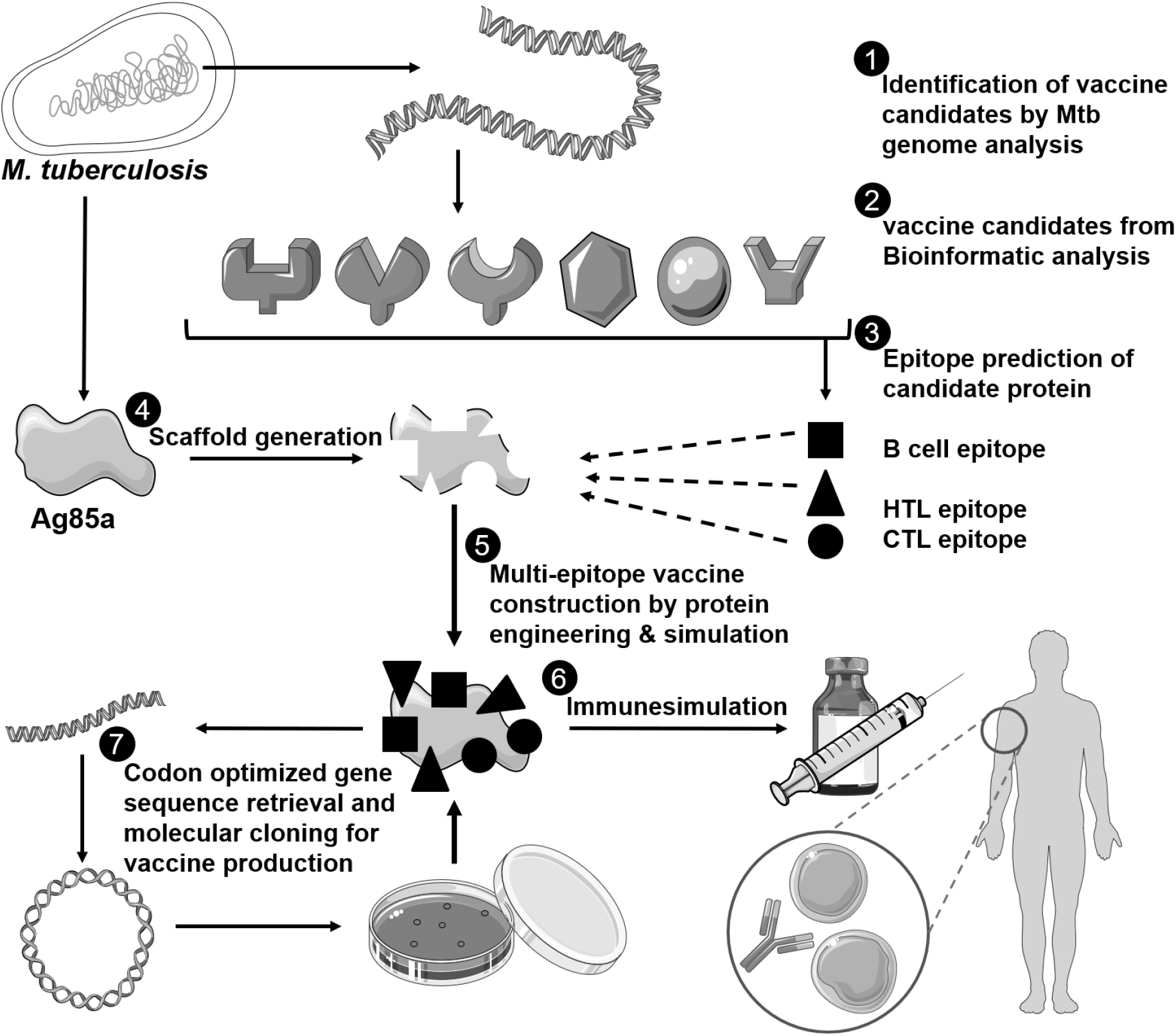
Graphical representation of the strategy to design scaffold based multiepitope vaccine against Mycobacterium tuberculosis.

## RESULTS AND DISCUSSION

### Retrieval of sequence and antigenicity of selected proteins

Mining the existing literature revealed nearly 100 secretory proteins that are involved in virulence or are essential for the survival of Mtb. While selecting potential vaccine candidates, only the secretory proteins that are either essential or responsible for virulence and show a >0.4 antigenicity score on VaxiJen 2.0 are selected (Table 1)[21]. The following proteins, RV0101, RV1161, RV3646c, RV1918c, RV3870 and RV3810 showing highest antigenicity score among this pool were selected for further analysis. Out of these six proteins two are responsible for virulence (RV0101, RV1161), three are essential genes (RV1918c, RV3646c, RV3810) and RV3870 is both virulent and essential.

**Table 1:**
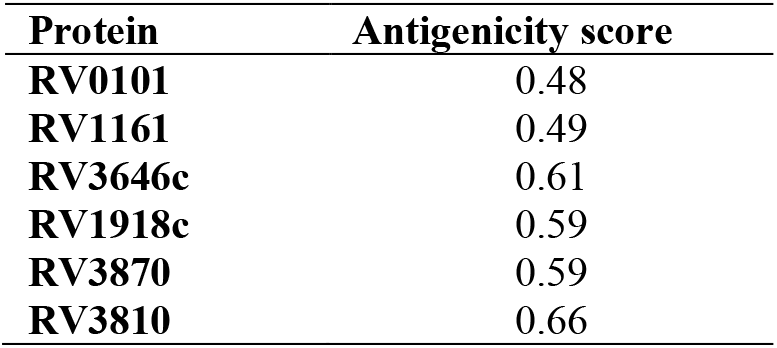
Antigenicity determination.

### Prediction of B-cell epitopes

The B-cell epitopes from the aforesaid proteins were predicted based on the output of ABCpred [22]. The peptides with a score above 0.85 were chosen for further analysis, a higher score indicates a high immunogenicity. From this pool, peptides which are neither allergic or toxic were selected for MEV construction. The sequence of the B-cell epitopes along with their antigenicity scores, allergenicity and toxicity predictions are tabulated in Table 2.

**Table 2:**
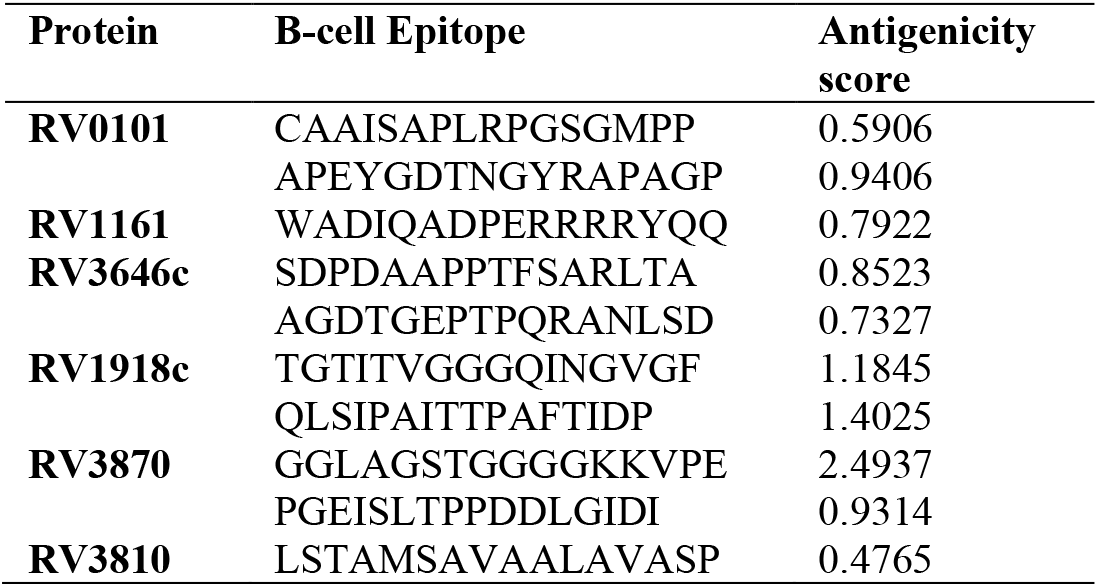
B-cell epitope prediction. **(all predicted epitopes are non-allergic and nontoxic)**

### Prediction of T-cell epitopes

Similarly, the above mentioned six proteins were further subjected to CTL (cytotoxic T lymphocyte) and HTL (helper T lymphocyte) epitope prediction. The selection of CTL and HTL epitopes were screened on the basis of MHC binding score, where lower score indicates higher affinity. From the selected T-cell epitopes, only the ones with high antigenicity (>0.4), and no allergenicity or toxicity were selected for further analysis. The IFN-γ production capability of these epitopes were also predicted and those peptides that induce IFN-γ were selected for MEV construction. The HTL, CTL epitopes along with their antigenicity score, allergenicity, and toxicity predictions are tabulated in Table 3 and 4.

**Table 3:**
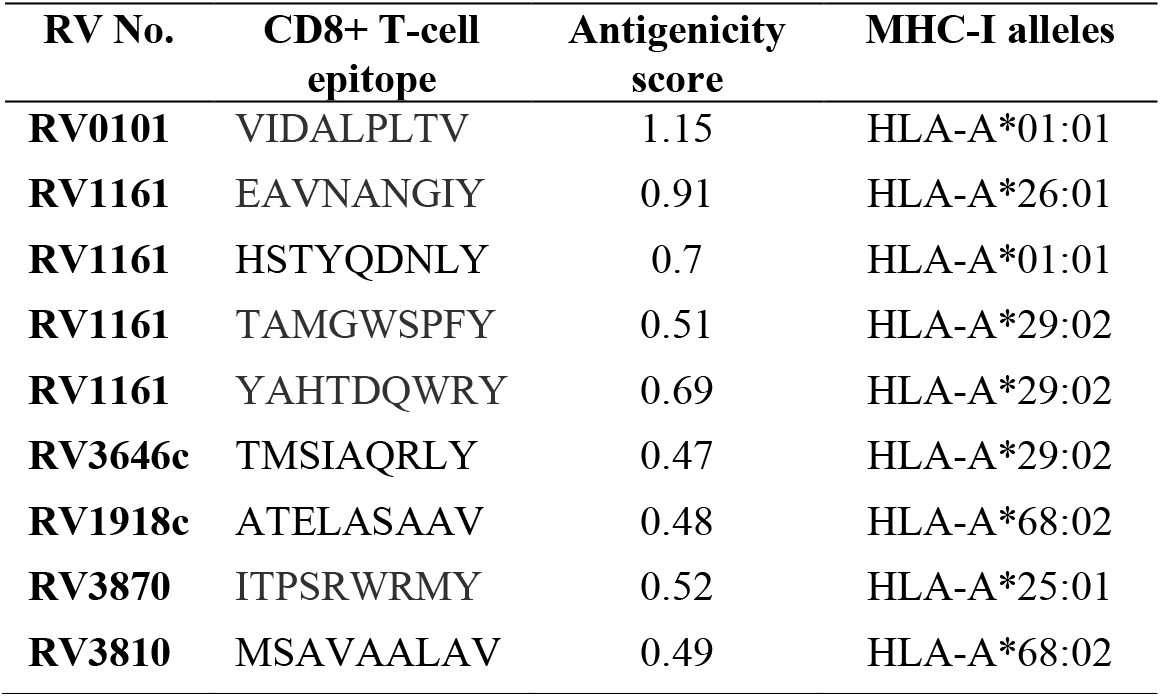
CD8+ T-cell epitope prediction. **(all predicted epitopes are non-allergic and nontoxic and induce IFN-γ)**

**Table 4:**
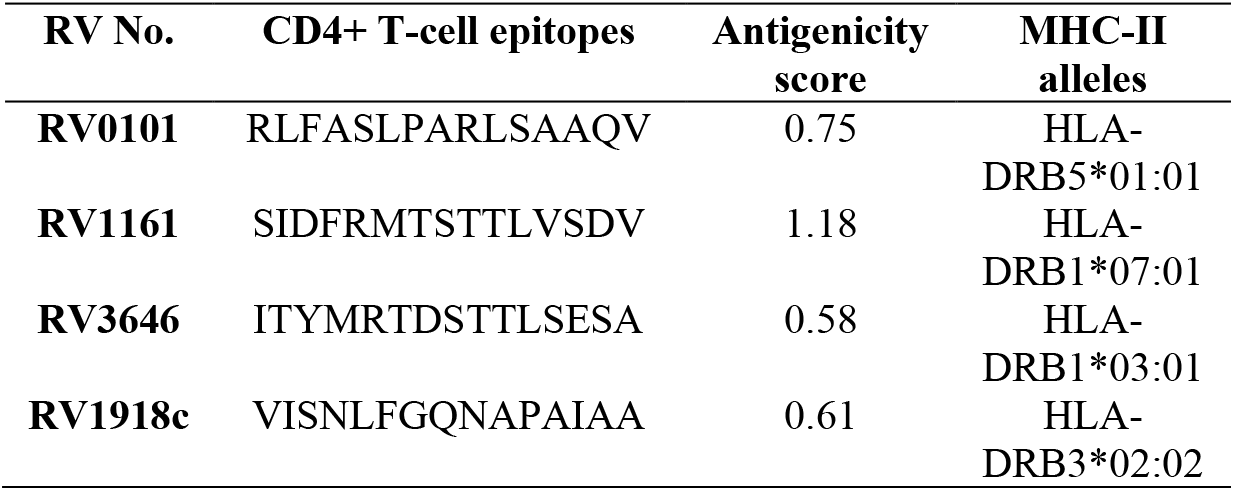
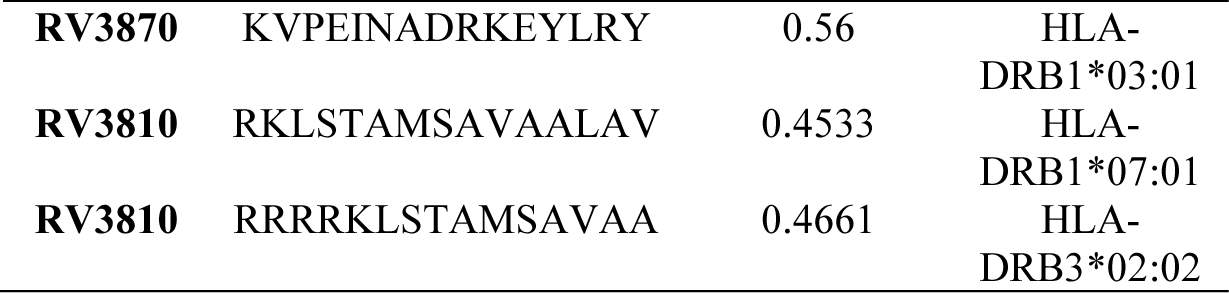
CD4+ T-cell epitope prediction. **(all predicted epitopes are non-allergic and nontoxic and induce IFN-γ)**

### Construction of Multi-Epitope Vaccine

To construct the scaffold based multi-epitope vaccine, the crystal structure of Ag85A protein of Mtb was used as scaffold. Initially the large loops of Ag85A were replaced by amino acid sequences of the predicted, B-cell, HTL and CTL epitopes. Several rounds of construction, deconstruction and successive cycles of model prediction using Robetta web tool provided the structural details of the selected epitopes (Table 5 and 6). Then the structural elements were strategically placed to fill the gaps in the truncated Ag85A protein which is serving as the scaffold. The truncated Ag85A is majorly harbouring the immunogenic part of the protein itself. Finally, to the N terminal part, a 45 amino acid long adjuvant, Human β-defensin-3, has been added. This newly constructed MEV, named as MEV6.2 consists of 605 amino acids. Figure 2A shows the 3D structure of the scaffold based multi-epitope vaccine MEV6.2 where different elements are colour coded, the Ramachandran Plot and distribution of secondary structure of this predicted model are presented in Supplementary Figure 1 and 2 respectively. Figure 2B depicts the deconstructed form of the MEV, following the same colour code, to provide a better understanding of the assemblage of the structural elements in the MEV6.2. The detailed sequence and structure of each epitope is described in Table 5 and 6. Beside MEV6.2, another vaccine has been constructed using the same epitopes, following the conventional approach, of multi-epitope vaccine construction where no scaffold has been used, we refer to it as MEV1. The predicted epitopes were simply glued one after another using linker along with the adjuvant at the N-terminus. The amino acid sequence of MEV6.2 and MEV1 are presented in a colour coded mode in Figure 3. The predicted 3D structure of MEV1 (503 amino acids) is illustrated in Figure 4, the Ramachandran Plot and secondary structures are presented in Supplementary Figure 3 and 4 respectively.

**Table 5:**
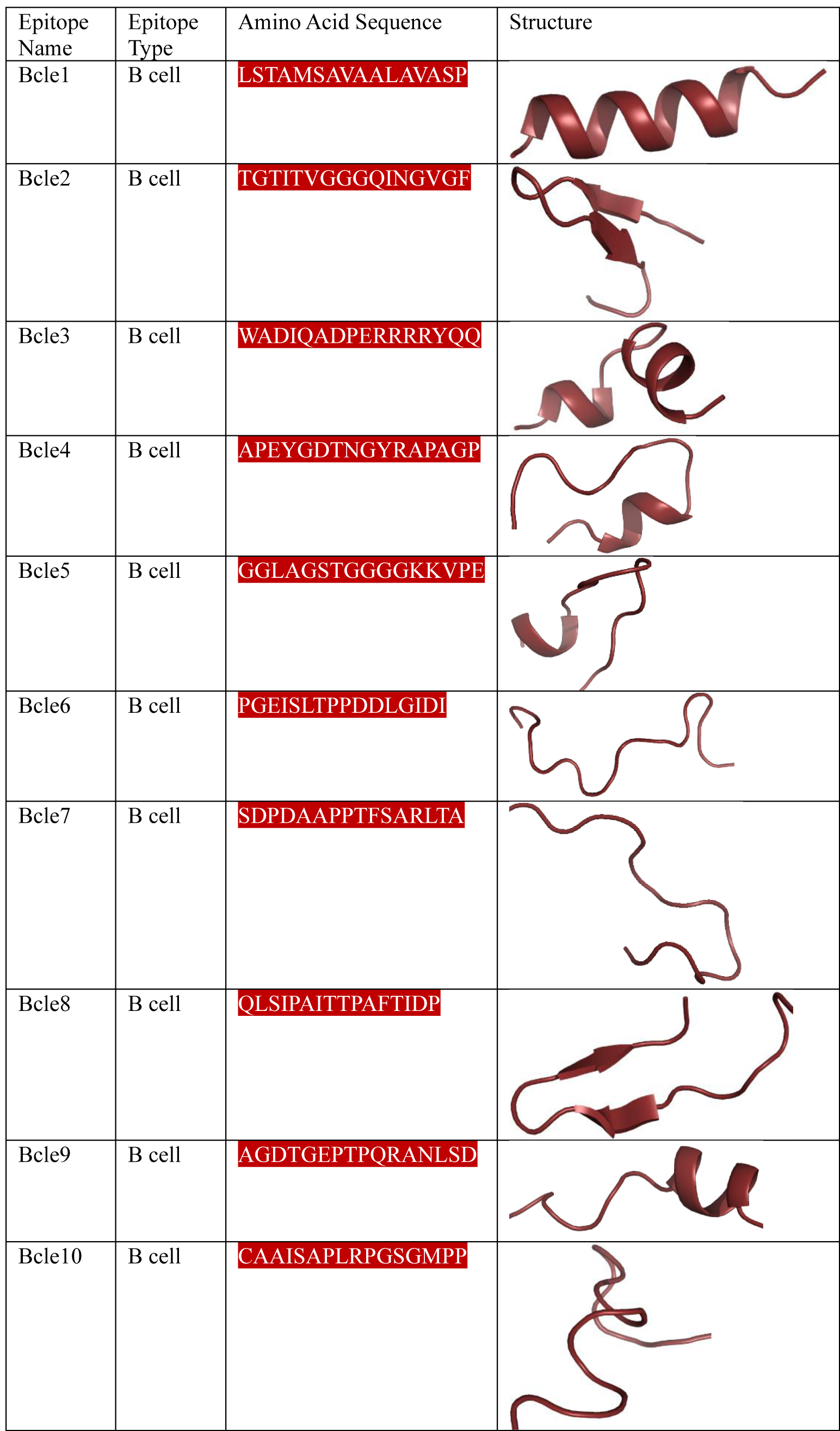
Predicted structures of B-cell epitopes.

**Table 6:**
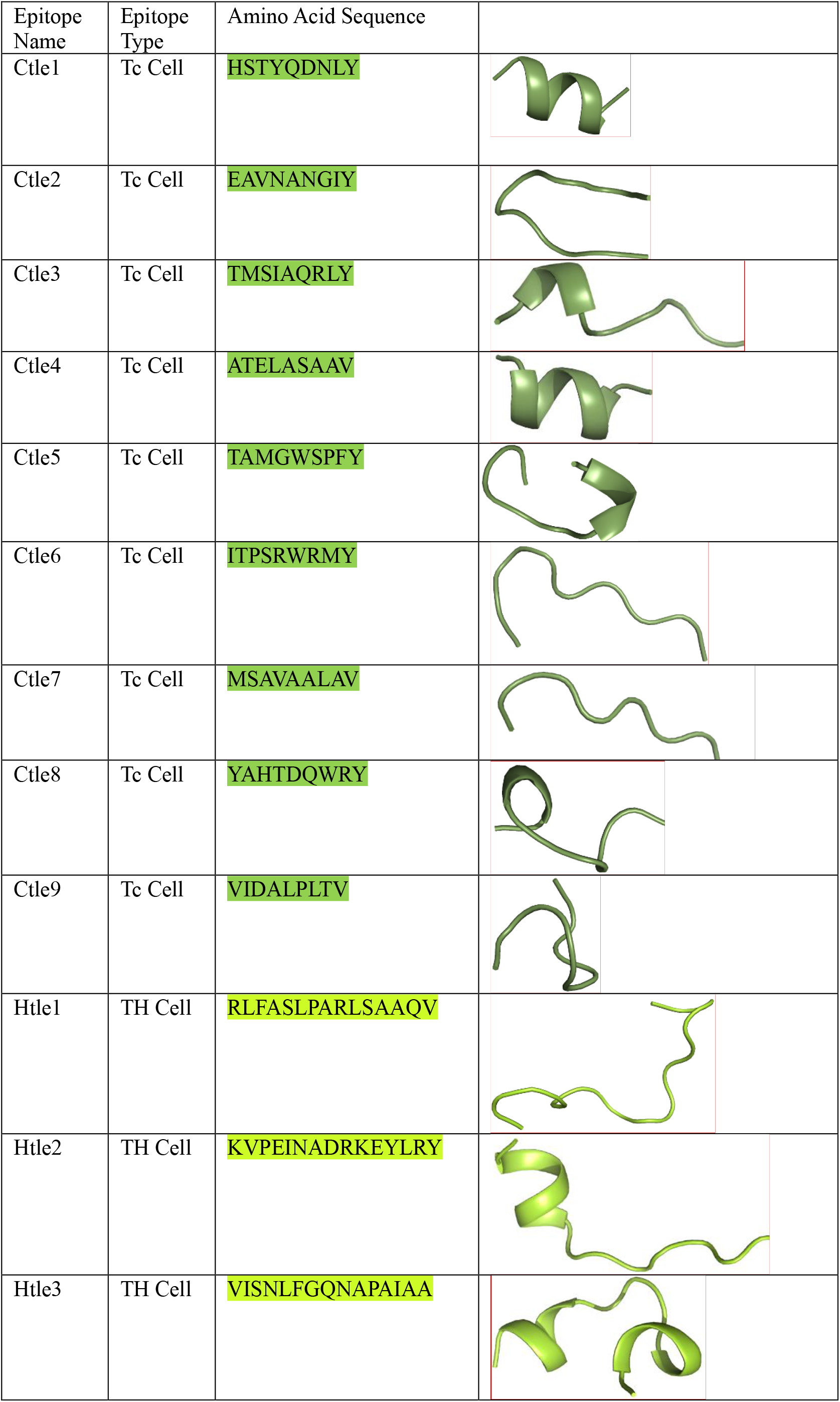
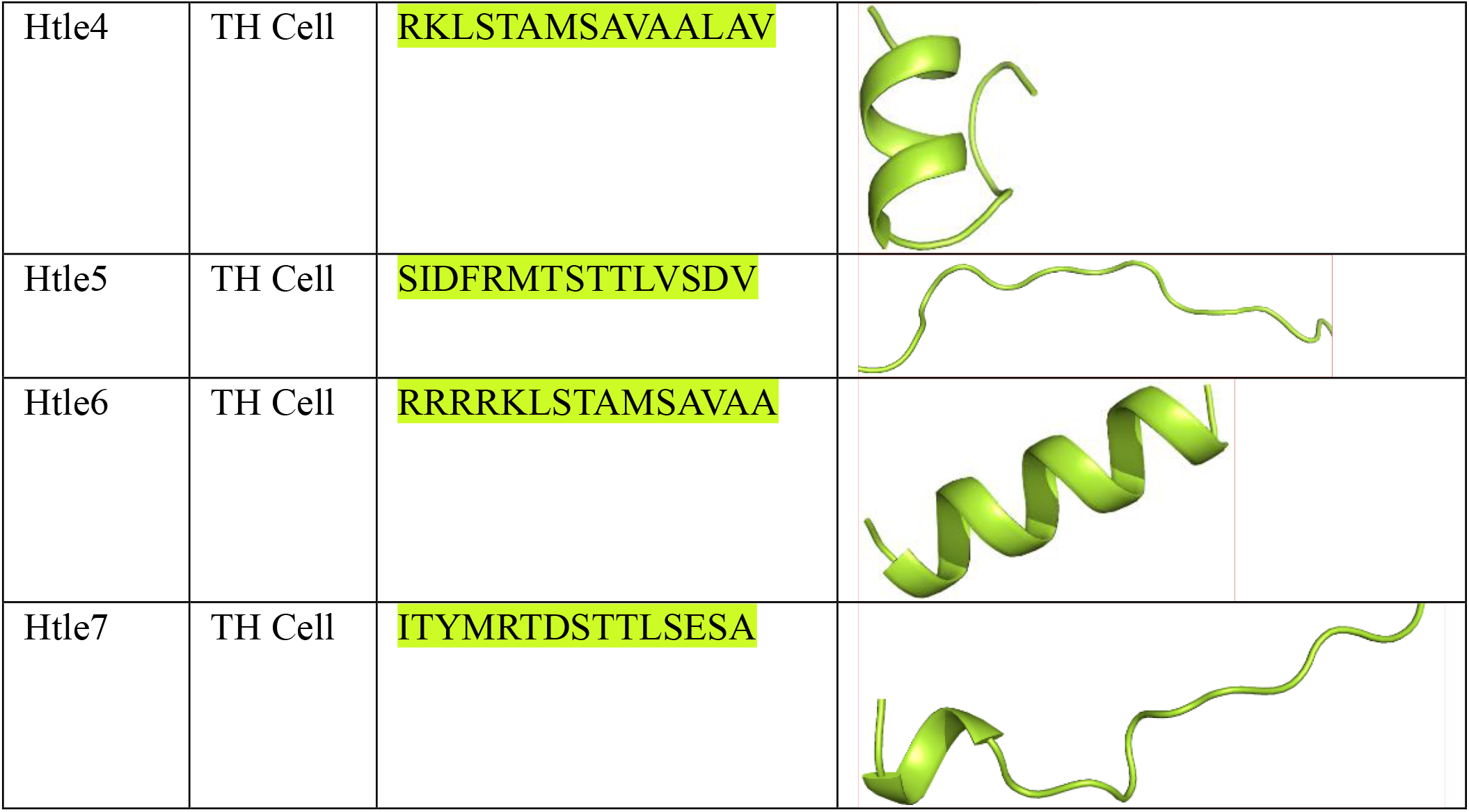
Predicted structures of T-cell epitopes.

**Figure 2:**
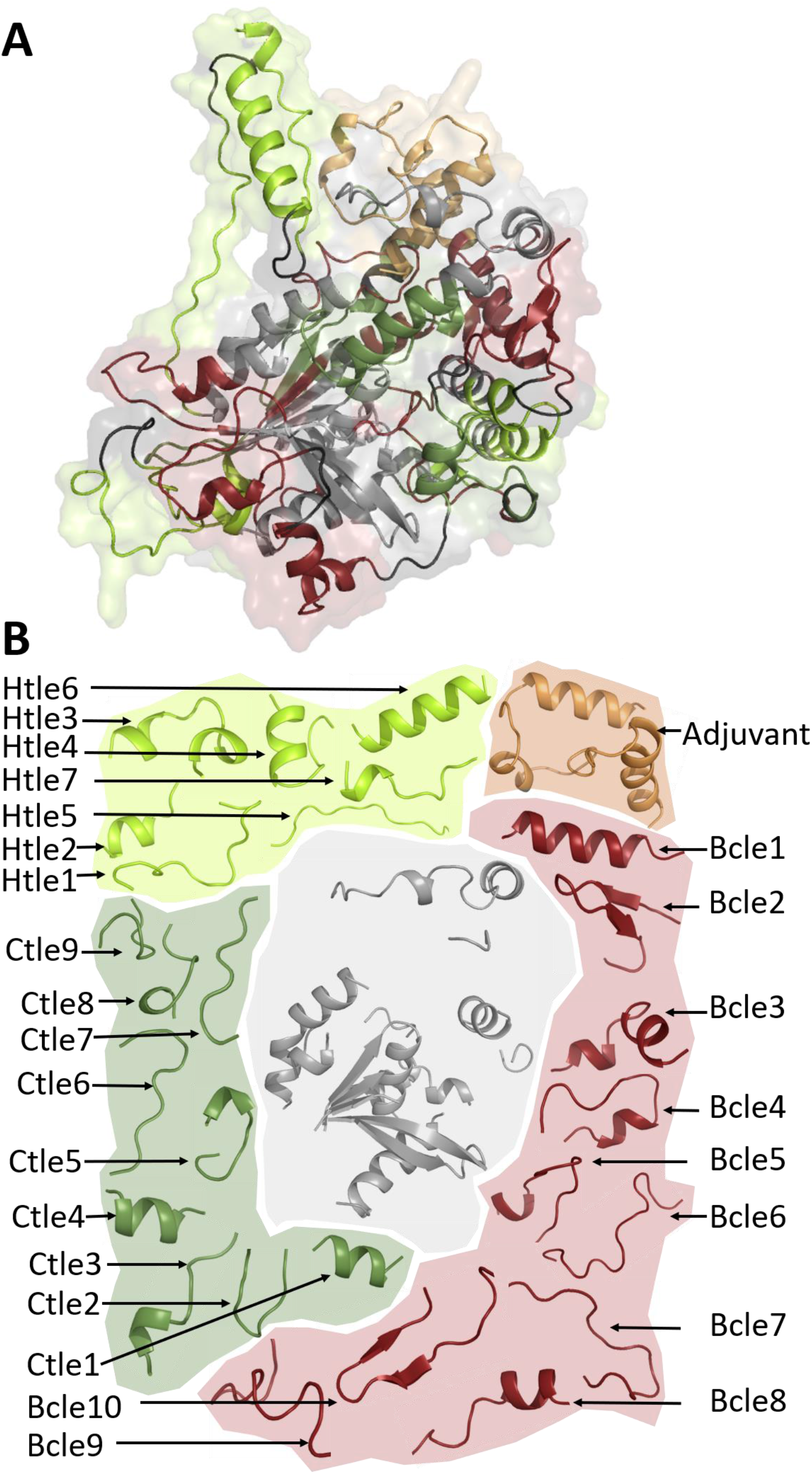
Structure of MEV6.2 in cartoon representation where B cell, cytotoxic T cell and Helper T cell epitopes are coloured as ruby, smudge and lemon respectively. The adjuvant is shown in light orange and the scaffold (Ag85A) protein in light grey (A). MEV6.2 structure has been deconstructed into its constituent B cell epitopes (Bcle1 – Bcle10), cytotoxic T cell epitope (Ctle1 – Ctle9), Helper T cell epitope (Htle1 – Htle7), scaffold (Ag85A) and adjuvant, presented in the identical colour code as described above. Each set of structural elements are within a shadowed boundary of the same colour.

**Figure 3:**
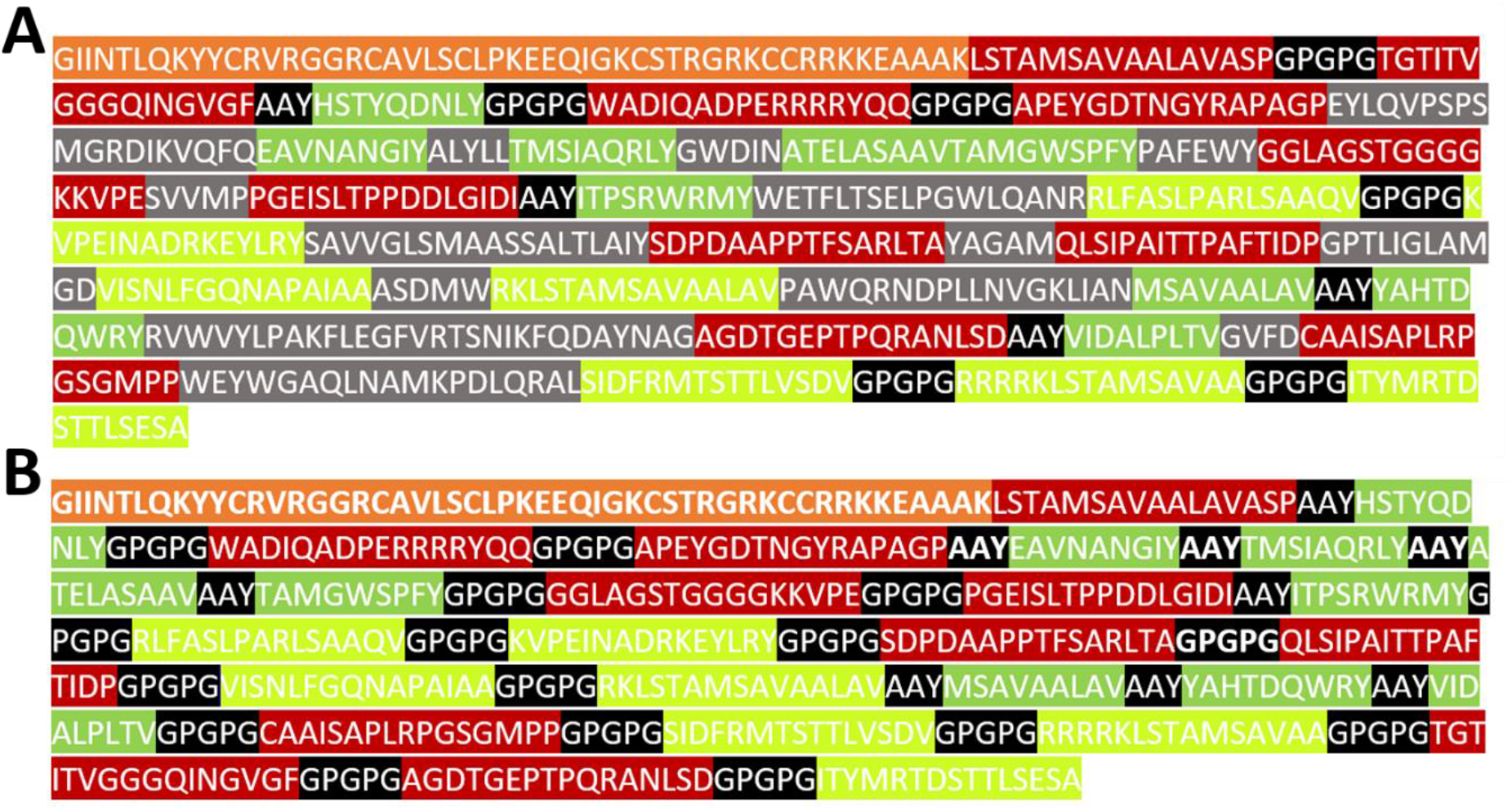
Amino acid sequence of MEV6.2 (A) and MEV1 (B). B cell, cytotoxic T cell and Helper T cell epitopes are coloured as ruby, smudge and lemon respectively. The adjuvant is shown in light orange and the linker are in black. In MEV 6.2, scaffold (Ag85A) protein sequence is shown in light grey.

**Figure 4:**
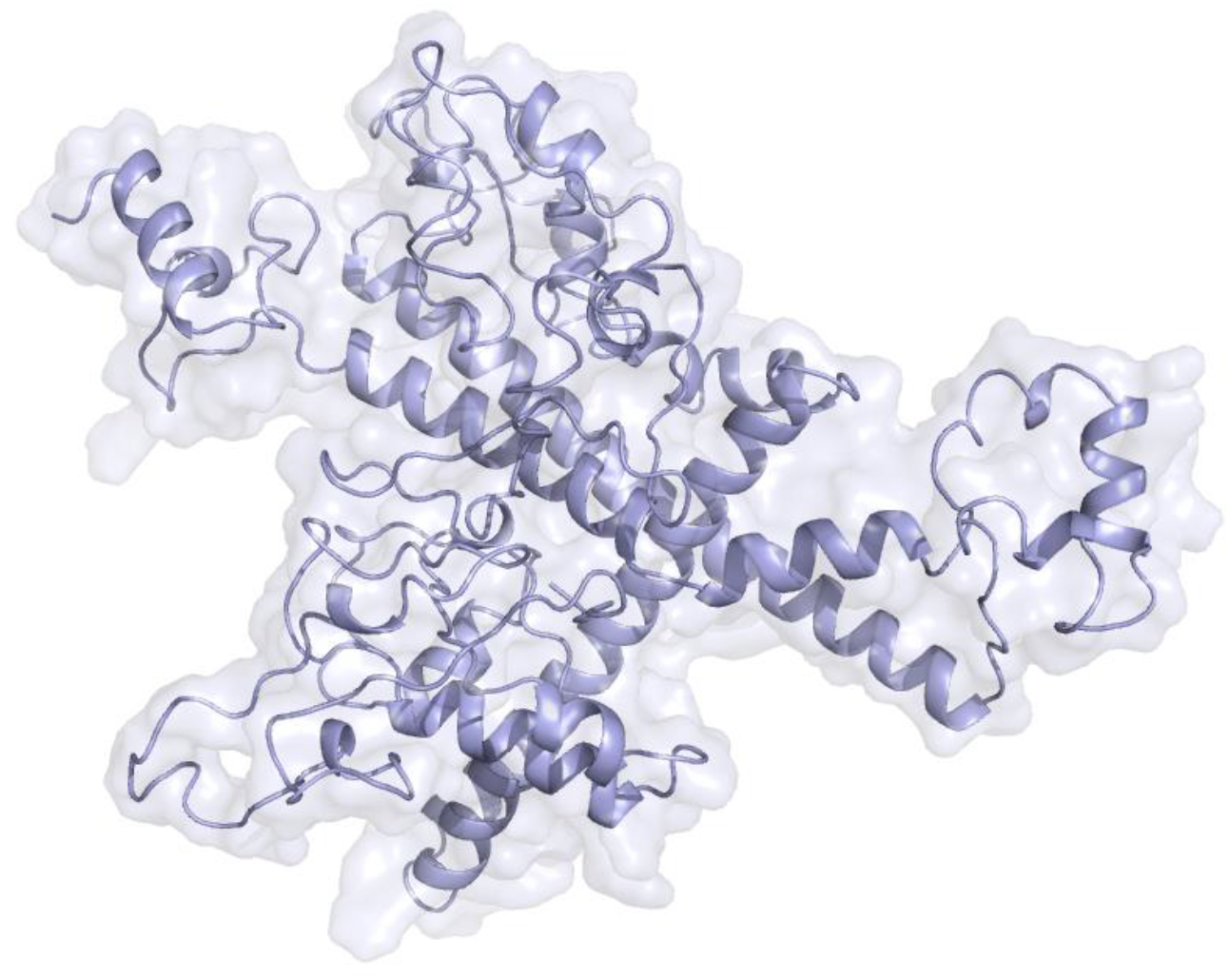
Tertiary structure of MEV1 presented in cartoon mode with an essence of surface.

### Proteasomal cleavage sites

Proteasomal cleavage and TAP binding predicts antigenicity of the constructed vaccine. For MEV6.2, NetChop [23] predicted 204 proteasomal cleavage site above threshold value (>0.5) (Supplementary Figure 5).

### Molecular dynamics simulation

Figure 5 describes the analysis of simulation trajectories of MEV. Figure 5A clearly depicts the RMSD (root mean square deviation) of MEV1 (average RMSD: 0.75nm) during the 100ns time scale is significantly greater than the RMSD of MEV6.2(average RMSD: 0.50nm), indicating MEV6.2 is much more stable over the course of simulation than MEV1, as expected from the tertiary structure of the proteins. Figure 5B compares, the Radius of gyration (Rg), which indicates compactness of the protein. The average Rg for MEV6.2 is 2.58nm which is lower than the Rg of MEV1, 2.73nm. A lower Rg value of MEV6.2 despite of its bigger size is indicative of its stability and compactness over the course of simulation. The residue wise root mean square fluctuation (RMSF) of MEV6.2 (Figure 5C) and MEV1 (Figure 5D) shows that MEV6.2 is much more stable (average RMSF: 0.17nm) compared to MEV1 (average RMSF: 0.30nm). The simulation data proves that the scaffold-based structure is more stable compared to the one designed with conventional approach. Hence MEV6.2 has been chosen for further analysis.

**Figure 5:**
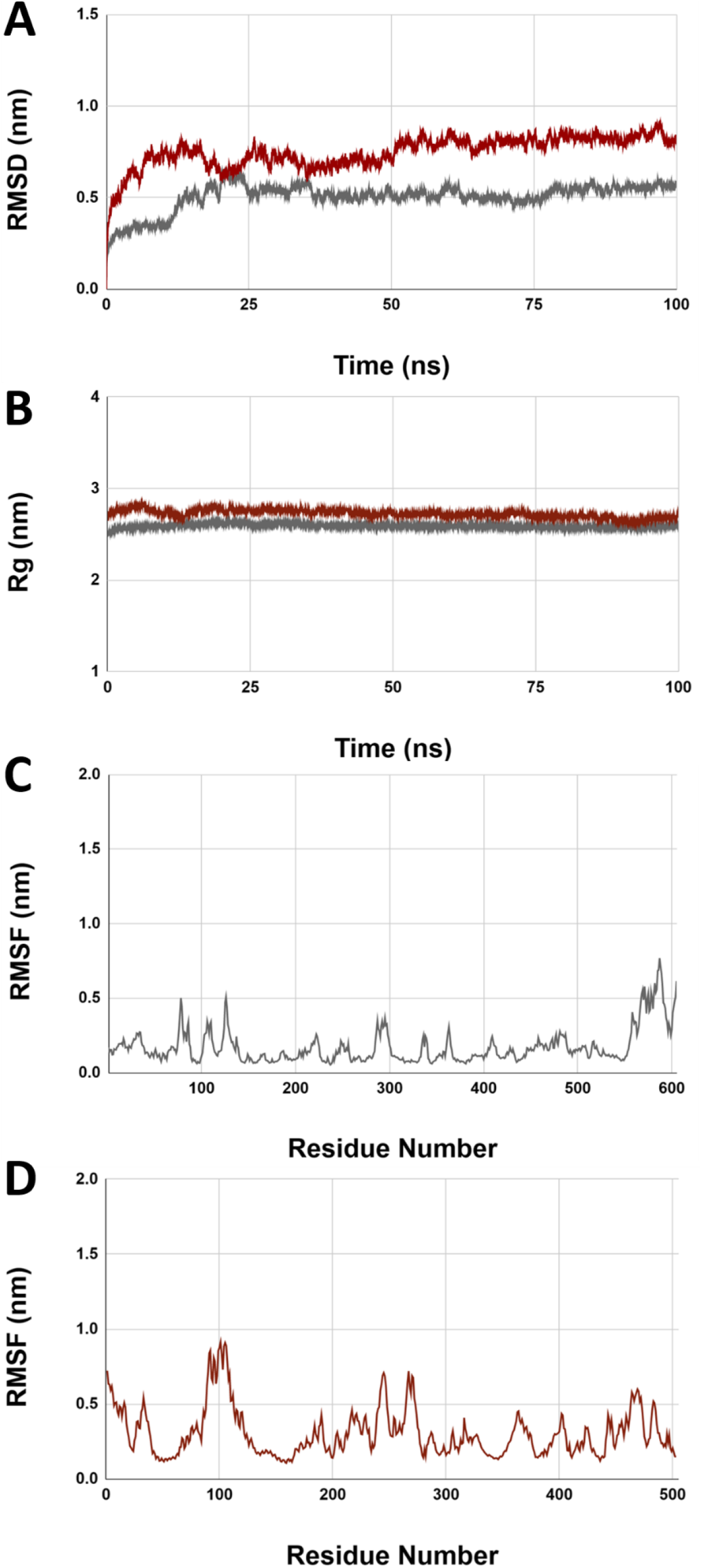
Graphical representation of molecular dynamics simulation trajectory analysis of MEV6.2 and MEV1 from 100ns time frame. (A) Comparison of RMSD (nm) of MEV6.2 (in light grey) and MEV1 (dark maroon). (B) Comparison of Rg (nm) of MEV6.2 (in light grey) and MEV1 (dark maroon). (C)RMSF (nm) of MEV6.2 (in light grey). (D) RMSF (nm) of MEV1 (dark maroon).

### Topology and structural flexibility

The detailed analysis of the tertiary structure of MEV6.2 is presented as the topology diagram in Figure 6A where it shows the protein is composed of 20 α helices, and 10 β strands connected by loops. The distribution of the secondary structure elements of the protein is presented in Supplementary Figure 2. All the β strands, majorly came from Ag85A scaffold whereas the α helices and connecting loops were contributed by the B-cell and T-cell epitopes (Table 5 and 6). Superposing the PDB structures extracted at 10ns intervals from the simulation trajectories indicates the major flexibility is coming from the loop regions and the N-terminal part, marked with black arrow heads in Figure 6B. The core structure remained stable, throughout the simulation, which is in good agreement with the RMSD and RMSF plots discussed earlier. The structural flexibility of MEV6.2 was also predicted using CABS-flex server with an average RMSF of 0.11nm (Supplementary Figure 6).

**Figure 6:**
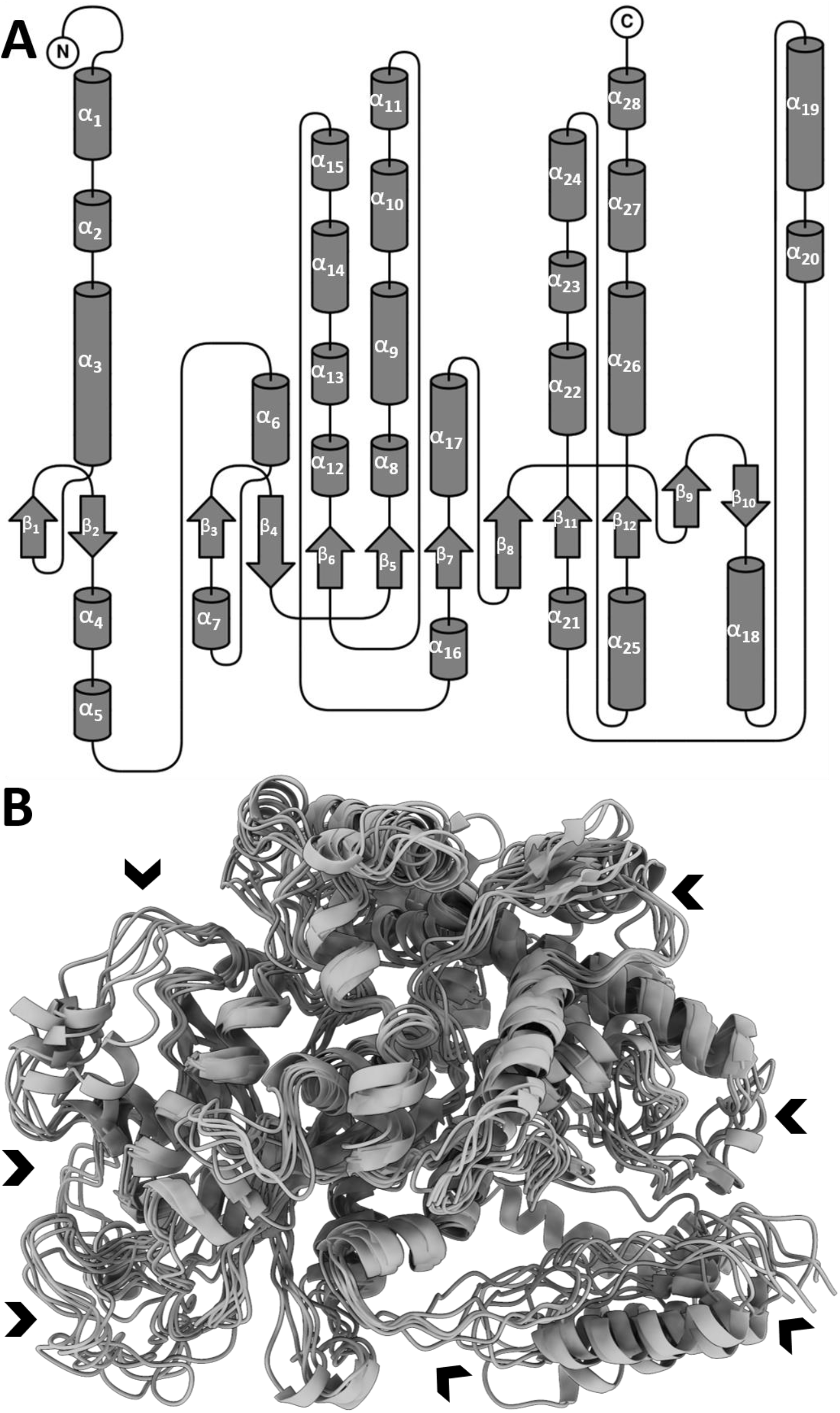
(A) Topology diagram of MEV6.2. The structure is composed of 10 β strands and 20 α helices. (B) Concerted motion derived from molecular dynamics simulation trajectories of MEV6.2. The Solid black arrow heads are indicating the flexibility of loop regions during the course of simulation.

### Interaction of MEV with TLR2

The docked complex of MEV6.2-TLR2, obtained from ClusPro server [24] is presented in Figure 7A. The binding free energy of MEV6.2-TLR2 was found to be -125.9kcal/mol, calculated using Molecular Mechanics/Generalized Born Surface Area (MM/GBSA) using HawkDock platform [25]. It has been found that the binding free energy for MEV1-TLR2 complex is -99.87 kcal/mol, suggesting MEV6.2 has a higher affinity for TLR2. The residues with major contribution, in MEV6.2-TLR2 complex formation, are tabulated in Figure 7B and C. From the MD simulation data, it has been found that MEV6.2-TLR2 forms a stable complex throughout the course of simulation (Supplementary Figure 7).

**Figure 7:**
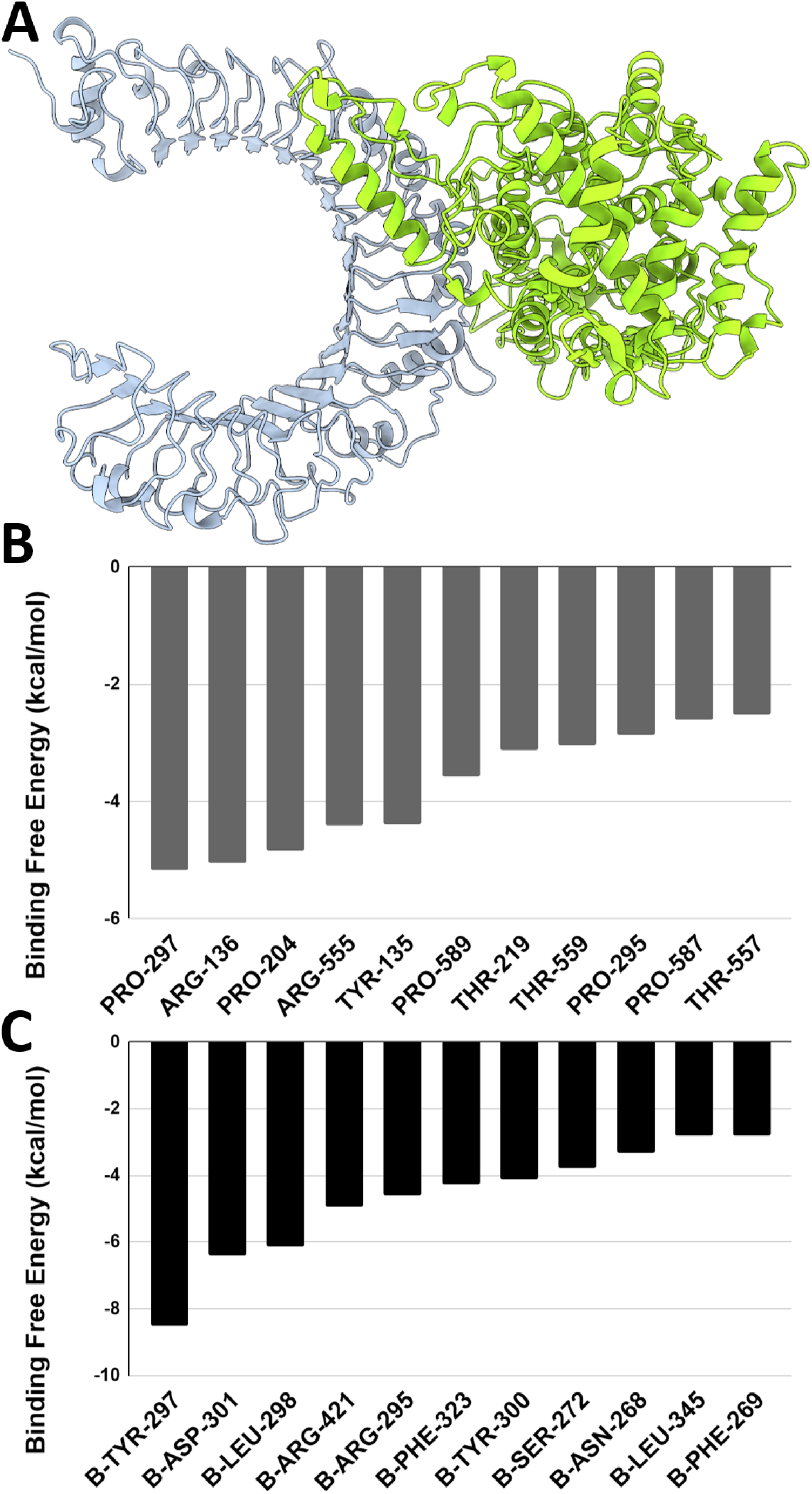
(A) Complex structure of MEV6.2 and TLR4 presented in cartoon, MEV6.2 in and TLR4 in. Amino acid residue wise binding free energy calculation: amino acid residues of MEV6.2 (B) and TLR4 (C) contributing majorly in complex formation.

### In-silico immune stimulation of MEV

The ability of MEV6.2 vaccine to induce B-cell, T-cell, Dendritic cell, Natural Killer cells and other immunomodulators have been predicted by C-IMMSIM server, presented in Figure 8. Production of IgG and IgM antibodies are boosted after the second and third injection. The concentration of B-cell population is increasing with each booster dose, with majority of the population belonging to active B-cell. The number of Plasma cells are also increasing with each booster dose, where the majority of plasma cells secrete IgM antibody after second booster dose and almost equal number of IgM and IgG antibodies after the third dose. There is almost threefold increase of Helper T cells after the second and third booster dose, the number of memory TH cells are highest at the end of tertiary booster dose. The number of Cytotoxic T cells and Natural killer cells is constantly high throughout the period of simulation. During the course of simulation, IFN-γ production is significantly high, other cytokines, TNF-b, IL10, IL4 are also stimulated.

**Figure 8:**
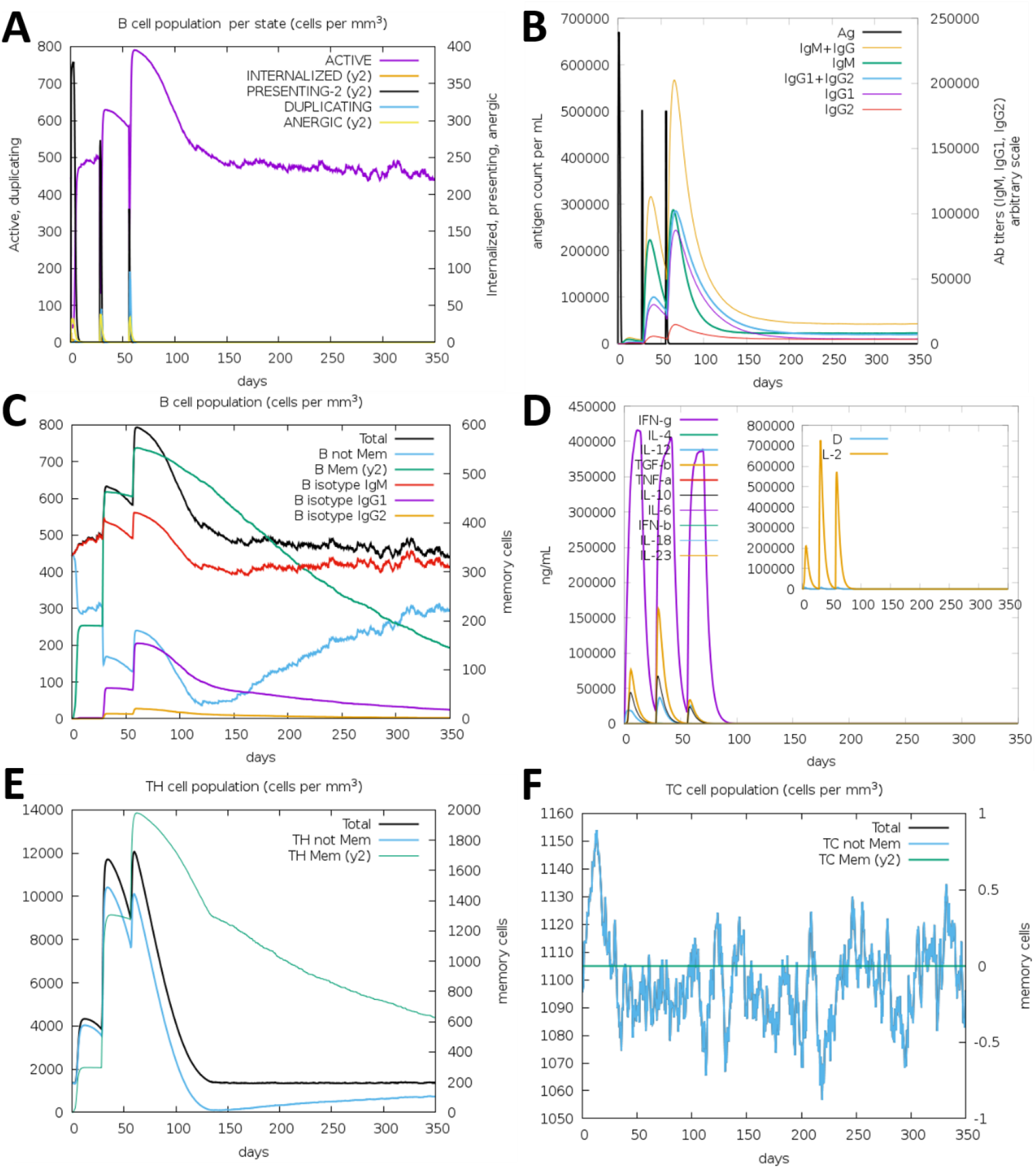
Immune simulation analysis: (A)B cell population per state, (B) IgG & IgM level increased after the administration of the vaccine, (C) Level of B cell population per mm^3^ in a window of 350 days, (D) Cytokines levels (elevated after the vaccine administrated, (E) TH cell population and (F) TC cell population in a window of 350 days.

### Codon optimization and in-silico construct design

As MEV6.2 has been found to be structurally stable, interacts strongly with TLR2, and immunogenic, hence it could be used as a potential vaccine against Mtb. For administration, the protein has to be produced in a highly pure form using recombinant DNA technology. The codon optimized DNA sequence deciphered from the amino acid sequence of MEV6.2 was inserted into an expression vector pET-28a using NdeI and XhoI restriction sites (Supplementary Figure 8). The codon optimized sequence in FASTA format is available in the Supplementary file, for *in-vitro* gene synthesis and cloning.

## CONCLUSION

A novel approach has been taken to construct a scaffold based Multi-Epitope Vaccine, against Mtb. The beauty of the method lies in using an under-trial vaccine candidate, Ag85A, as a scaffold to harbour potential B-cell and T-cell epitopes derived from computational immunology. The immunogenicity of the constructed MEV6.2 has been found to be higher than the scaffold protein alone and has the potential to elicit both B-cell and T-cell mediated immune response. Several steps of validation were adapted to ensure that MEV6.2 is structurally more stable than the MEV created by conventional approach. The tertiary structure of MEV6.2 shows a compact globular shape, that did not scramble during the course of simulation. Which makes it more suitable for large scale protein production, purification and storage.

MEV construction has become a lucrative area of research, where the conventional approach is to join the epitopes one after another with suitable linker. This strategy produces a polypeptide which is immunogenic and may show good result in immune simulation due to the presence of carefully selected B-cell and T-cell epitopes. However, if we consider the structure, the MEV is mostly comprised of loops and a few helices, due to the lack of secondary structure elements like β sheets, the resulting MEV is less compact, and shows higher flexibility that may affect the stability of the MEV. Using Protein-engineering, to accommodate the chosen B-cell and T-cell epitopes in a cautiously organised scaffold can keep a balance between the different secondary structure elements and produce a compact tertiary structure with greater stability, as we have found in this study. It is an example, of designing MEV, that should be helpful to achieve a subunit-based vaccine, against Mtb. Similar method could be applied to design vaccines against other pathogens. The scaffold based, MEV6.2 could be helpful in eradicating tuberculosis, although it requires experimental validation.

## MATERIALS AND METHODS

### Antigenicity prediction

The antigenic proteins were selected through extensive literature mining. Sequences of all the antigenic proteins were retrieved from Mycobrowser database (https://mycobrowser.epfl.ch/). The antigenicity of these proteins was predicted using VaxiJen 2.0 server (http://www.ddg-pharmfac.net/vaxijen/) [21] where proteins with an antigenicity score of >0.4 were selected for further analysis.

### Prediction of B-cell epitopes

B-cell epitopes are predicted using ABCpred server (https://webs.iiitd.edu.in/raghava/abcpred/index.html) [22] with a default cut-off score of 0.51. The peptides selected in this step were further screened for antigenicity, allergenicity and toxicity prediction using VaxiJen, AllerTOP server (https://www.ddg-pharmfac.net/AllerTOP/) [26] and ToxinPred server (https://webs.iiitd.edu.in/raghava/toxinpred/algo.php)[27, 28] respectively.

### Prediction of T-cell epitopes

The nonameric Cytotoxic T-lymphocyte (CTL) epitope prediction was performed using NetCTL server (https://services.healthtech.dtu.dk/services/NetCTL-1.2/ ) [29]. For prediction of binding affinity, proteasomal C terminal cleavage and TAP (transporter associated antigen processing) cut-off values of 0.05,0.15 and 0.75 were set respectively. Epitopes selected on this step were additionally subjected to MHC-I allele binding ability using IEDB MHC-I binding prediction server (http://tools.immuneepitope.org/mhci/)[30, 31]. Helper T-lymphocyte (HTL) epitopes were predicted using IEDB MHC-II server (http://tools.iedb.org/mhcii/)[32] .

### Antigenicity, allergenicity and toxicity prediction

Antigenicity, allergenicity and toxicity of predicted epitopes were determined using VaxiJen server, AllerTop server and ToxinPred server respectively. The interferon-γ (IFN-γ) inducing capability of the T-cell epitopes were predicted using IFNepitope tool (https://crdd.osdd.net/raghava/ifnepitope) [33].

### Construction of multi-epitope vaccine

Two different vaccine constructs were designed, the conventional MEV1 and an engineered vaccine MEV6.2. *Construction of MEV1:* the selected B-cell, HTL and CTL epitopes were linked using suitable linkers as described by Das group [3], [34]. As an adjuvant, Human β-defensin-3 (45amino acids), was used as adjuvant and it was linked to the N-terminus by EAAAK linker [3]. The final construct was subjected to antigenicity prediction using Vaxijen 2.0 and its antigenicity and allergenicity was also predicted following similar method as mentioned before.

*Construction of MEV6*.*2:* to improve stability of the newly designed vaccine, the selected B-cell, HTL, CTL epitopes were positioned on a carefully designed scaffold protein. Ag85A of *Mycobacterium tuberculosis* was chosen to be the scaffold since it is a well-established vaccine candidate. Ag85A was truncated in such a way where the flexible non-immunogenic parts of the protein were removed and the selected epitopes were strategically placed to fill these gaps. This process was iterated by several cycles of construction, deconstruction and model building to obtain a compact globular tertiary structure. Finally, the MEV version 6.2 (here referred to as MEV6.2) was selected for further analysis. Once the scaffold based MEV was constructed, the Human β-defensin-3 protein has been linked to the N-terminus via EAAAK linker as an adjuvant. The epitopes were joined using similar linker as MEV1.

### Prediction of 3D structure

Tertiary structures of both MEV1 and MEV6.2 was predicted by Robetta (https://robetta.bakerlab.org)[35]. The predicted structures were further validated by PROCHECK (https://www.ebi.ac.uk/thornton-srv/software/PROCHECK/) [36].

### Molecular Dynamics Simulation

MEV6.2 and MEV1, both were subjected to 100ns MD simulation was performed using Gromacs 5.1.2 platform (https://www.gromacs.org/ ) [37]. The protein molecules were simulated using CHARMM27 forcefield[38], following the protocol described previously[39, 40].

### Prediction of proteasomal cleavage sites and flexibility analysis

Proteasomal cleavage was predicted using the NetChop server [23]. Flexibility of the MEVs was evaluated using CABS-Flex 2.0 server [41] selecting 50 cycles and 50 cycles between trajectories.

### Interaction of MEV with TLR2

Toll Like Receptor 2 (TLR2) has a significant role in *Mycobacterium tuberculosis* infection, therefore the interaction between TLR2 and the designed vaccines have been assessed by docking the vaccines with TLR2 using the ClusPro webserver (https://cluspro.bu.edu/ ) [24]. The binding free energy was calculated by MM/GBSA method using HawkDock (http://cadd.zju.edu.cn/hawkdock/) [25].

### Molecular dynamics simulation of MEV6.2-TLR2 complex

The stability of MEV6.2-TLR2 complex, was assessed by (MD) simulation. MD simulation trajectories were centred and corrected for to calculate, RMSD (Root Mean Square Deviation)[42].

### Immune Simulation of MEV6.2

C-IMMSIM web server (https://kraken.iac.rm.cnr.it/C-IMMSIM/ ) [43] was used for prediction of immunogenic profiles of humoral and cell mediated response for the MEV6.2. The predictions were made based on administration of 3 consecutive injections with intervals of 4 weeks with time steps of 1, 84 and 168 time steps (where 1 time step is equal to 8hours). Each dose contained 1000 vaccine particles and the entire simulation was performed for 350 days (1050 time steps).

### Codon optimization and in-silico cloning

The DNA sequence of the multi-epitope vaccine, was obtained using Reverse Translate tool (https://www.bioinformatics.org/sms2/rev_trans.html ) The obtained sequence was further codon optimized for expression in E. coli (strain K12) host using Java codon Adaptation (JCat) server (http://www.jcat.de/ ) [44], choosing ‘avoid the rho-independent transcription termination’, and ‘avoid prokaryote ribosome binding site’ options. In the output a >0.8 Codon Adaptation Index (CAI) is considered a good score and a GC content 30-70% is ideal. For cloning, plasmid vector pET-28a(+) and restriction enzymes, NdeI and HindIII were selected. The feasibility of the construct was verified using SnapGene software (https://www.snapgene.com).

## AUTHOR CONTRIBUTION

DH and AR conceptualized the project, designed the experiments, analysed the data, designed the MEVs, prepared the figures and wrote the manuscript. TM performed epitope prediction, modelling, docking and immune simulation. SR performed MD simulation. AKD provided the expertise and infrastructure for MD simulation.

