## supplemental tables and figures for "Designing a Novel 3D Scaffold for Multiepitope Vaccine Development: Engineering Ag85a Protein for Enhanced Stability and Antigenicity"

#### Supplementary Figure 1

**Ramachandran Plot of MEV6.2**

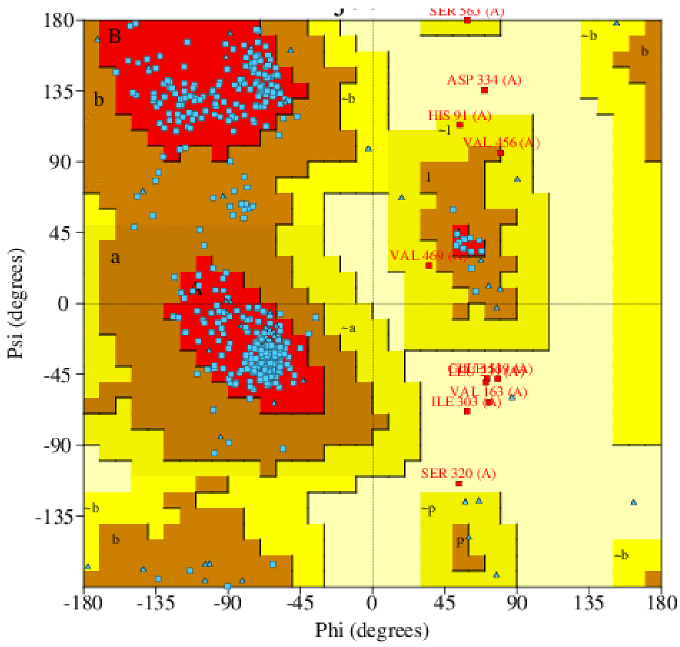


90.7% residues are in most favoured region, 7.1% residues are in additionally allowed region and 0.8% residues are in generously allowed region for MEV6.2

### Supplementary Figure 2

#### Secondary Structure of MEV6.2

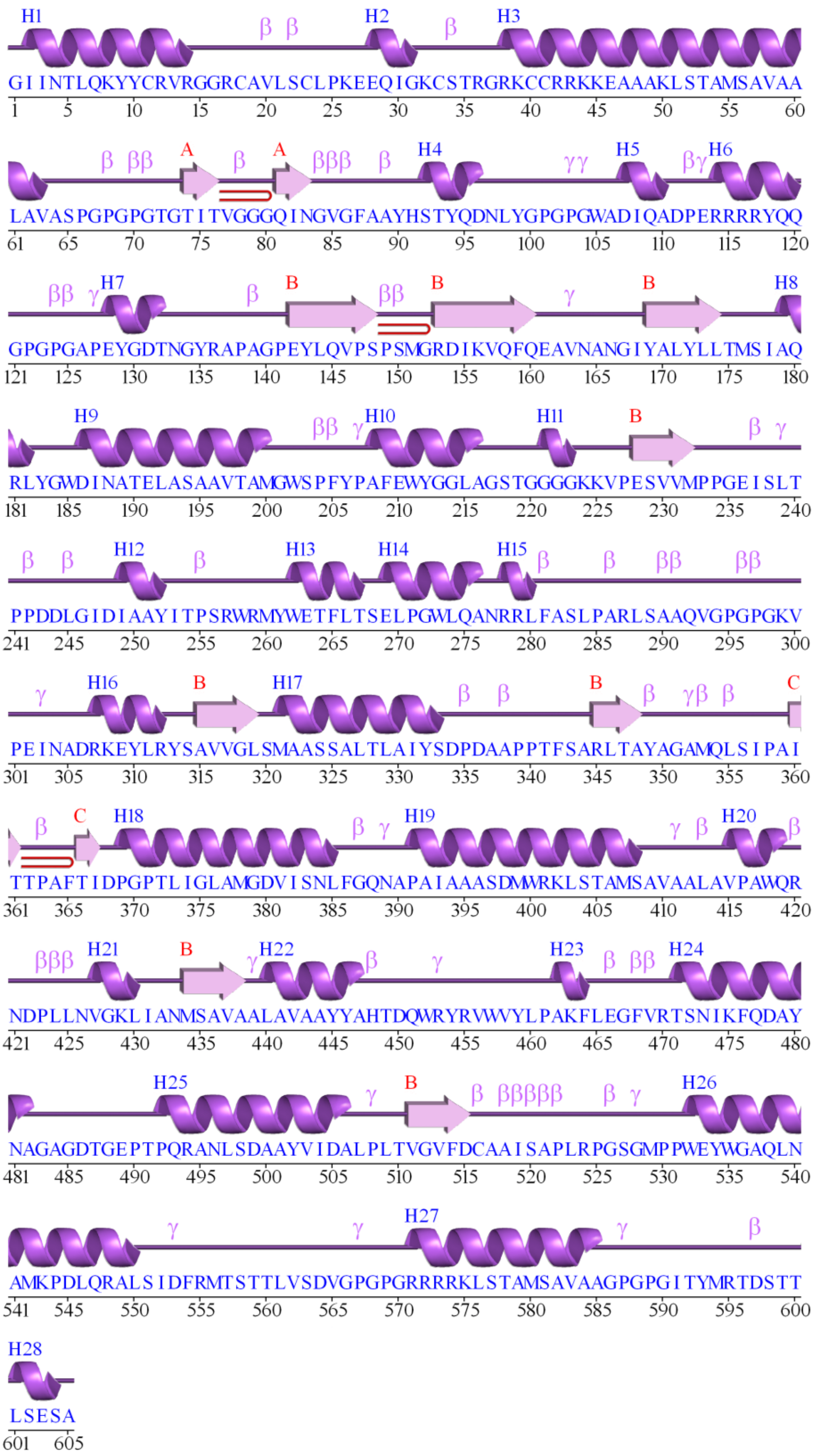

### Supplementary Figure 3

#### Ramachandran Plot of MEV 1

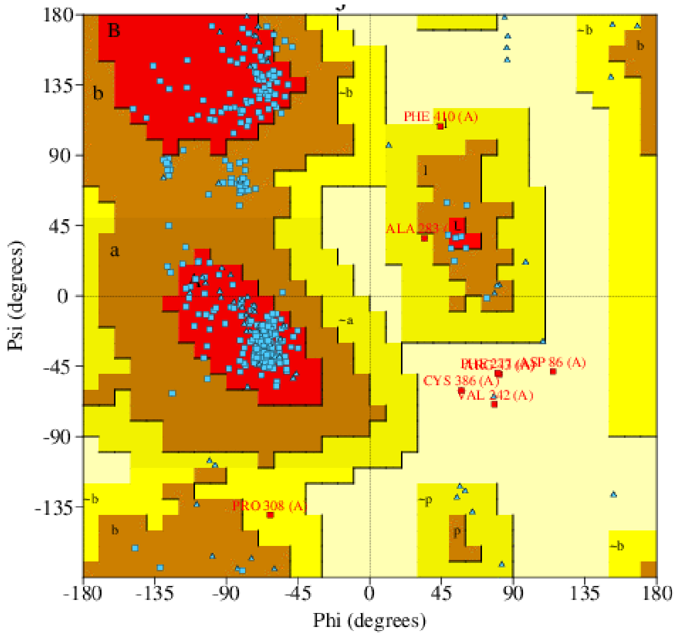


86.3% residues are in most favoured region, 11.8% residues are in additionally allowed region and 1.4% residues are in generously allowed region for MEV1

### Supplementary Figure 4

#### Secondary Structure of MEV1

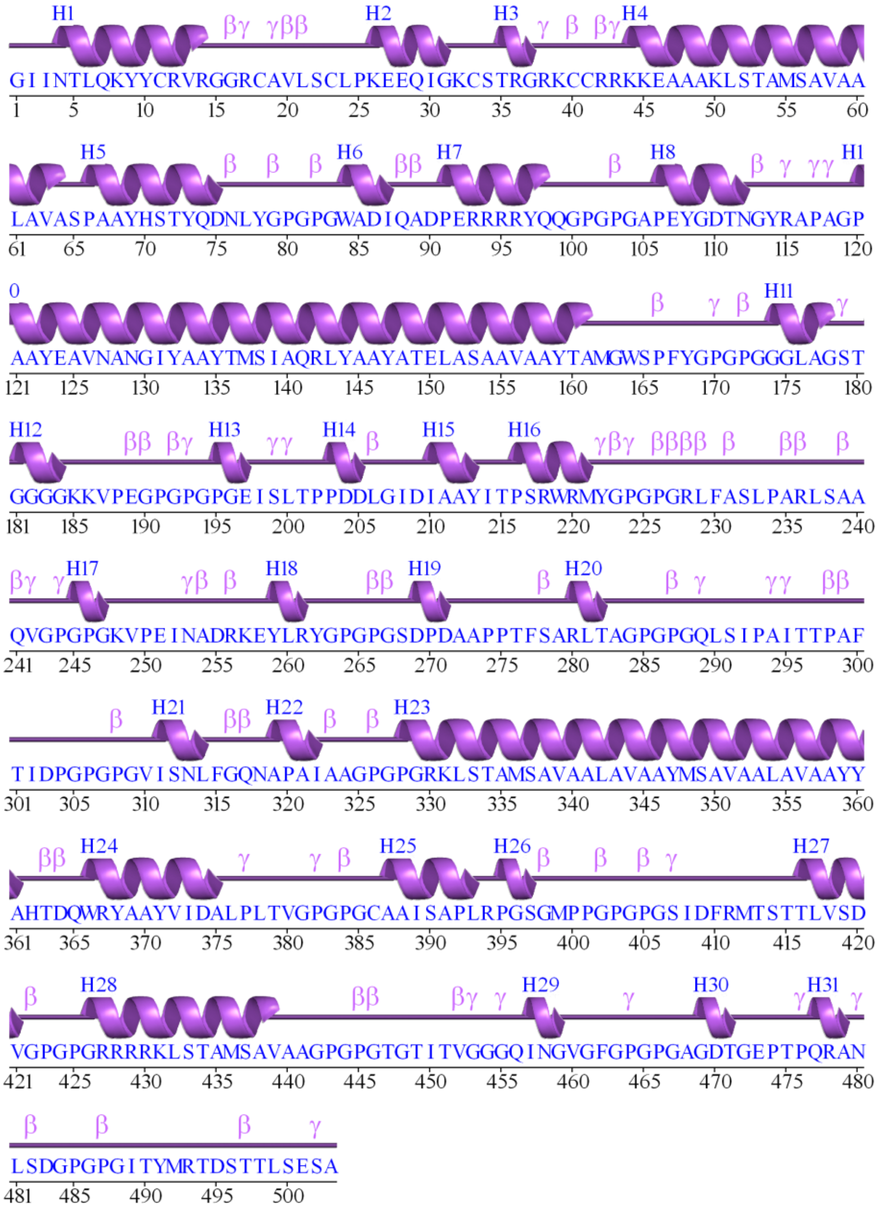

**Supplementary Figure 5**

**Prediction of proteosome cleavage site of MEV6.2**

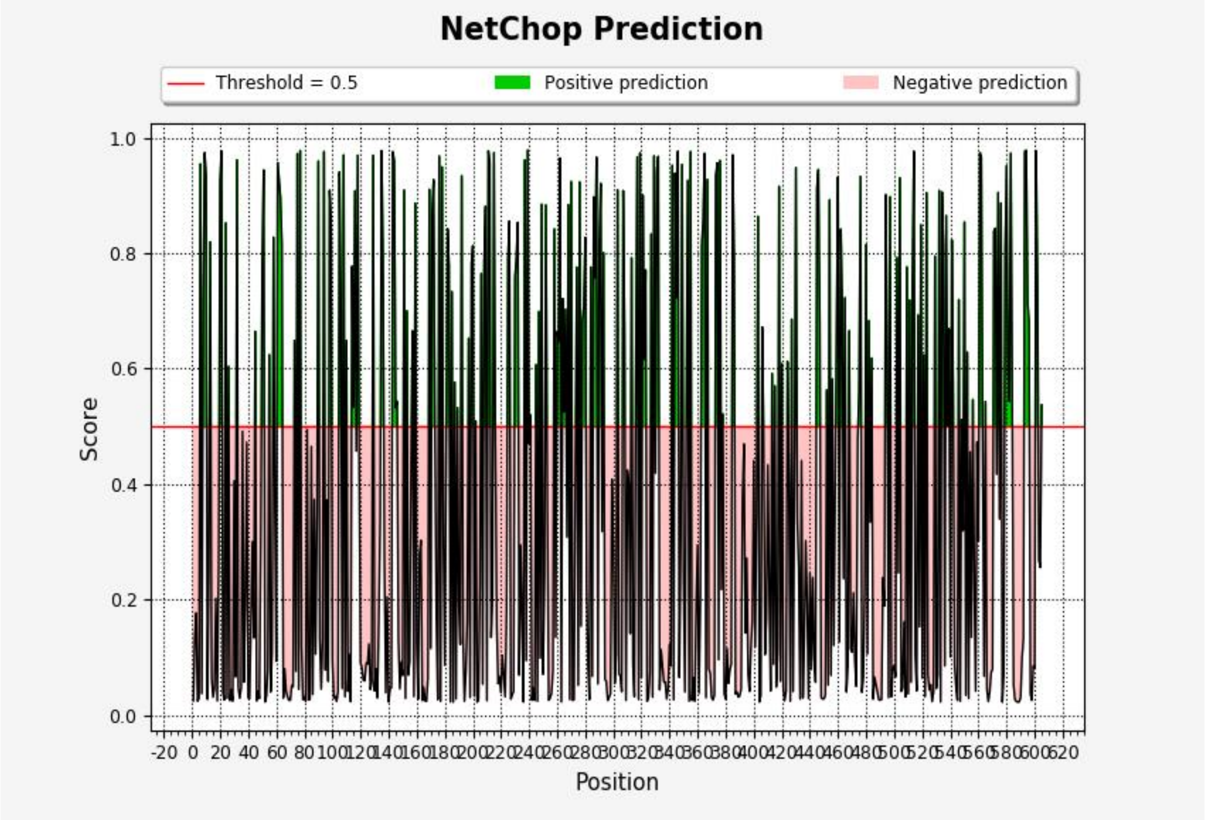

**Supplementary Figure 6**

**Structural flexibility of MEV6.2 predicted by CABS-Flex 2.0**

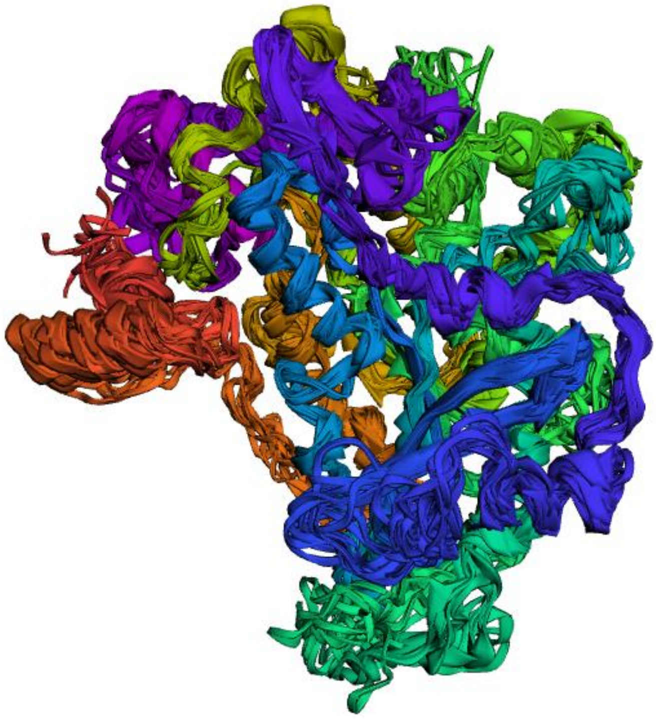

**Supplementary Figure 7**

**Graphical representation of molecular dynamics simulation trajectory analysis (RMSD in nm) of MEV6.2-TLR4 complex. Average deviation is 0.563nm.**

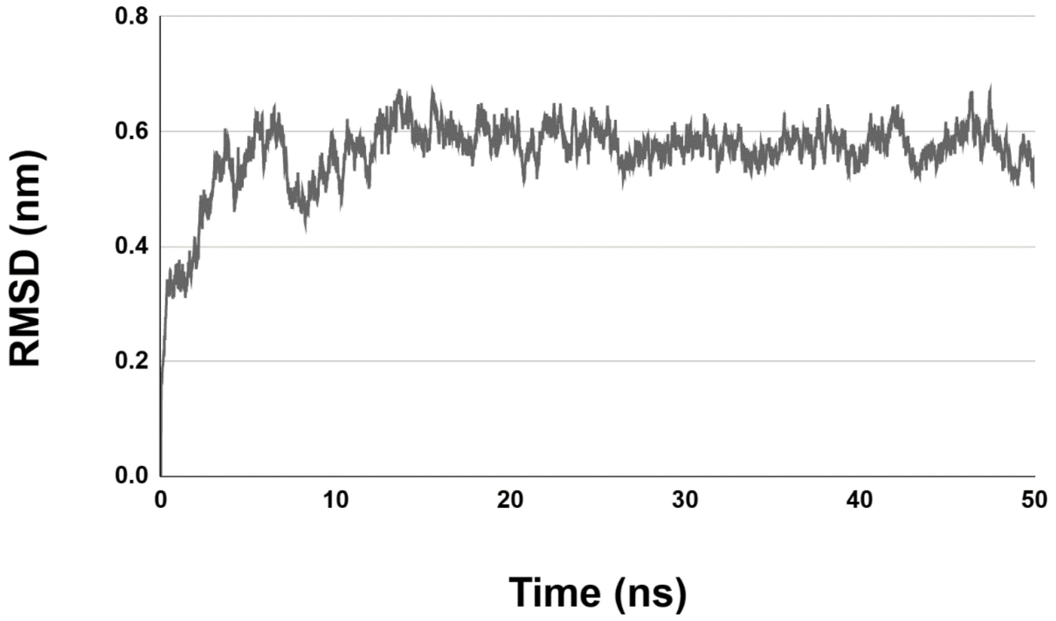

### Supplementary Figure 8

**MEV6.2, cloned in pET28a by Nde1 and Xho1 site. The construct is shown in red.**

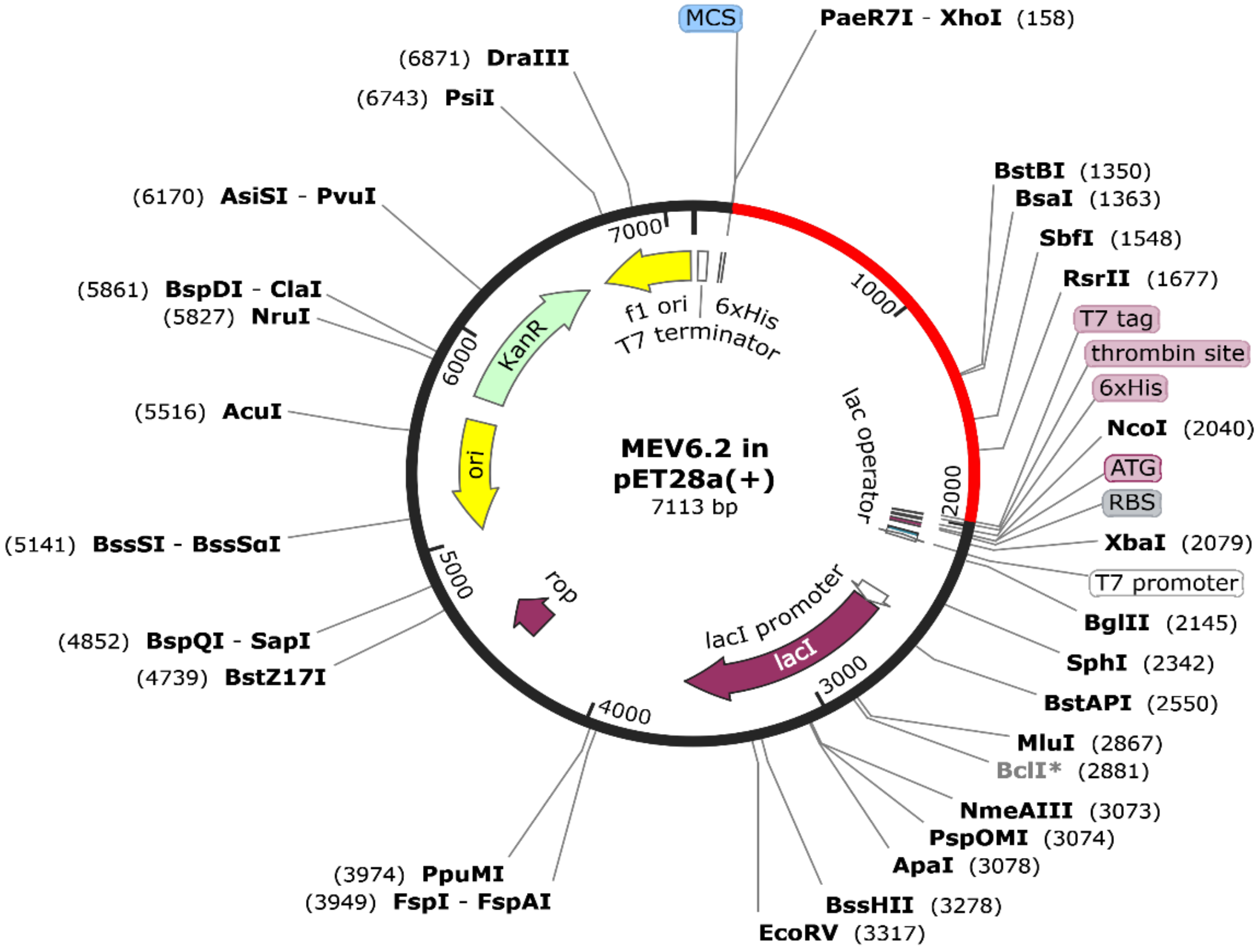

### MEV6.2 codon optimized sequence with NdeI and XhoI restriction site.

```
>MEV6_2_ARCDH_03062023
ATATGGGTATCATCAACACCCTGCAGAAATACTACTGCCGTGTTCGTGGTGGTCGTTGCG
CTGTTCTGTCTTGCCTGCCGAAAGAAGAACAGATCGGTAAATGCTCTACCCGTGGTCGTA
AATGCTGCCGTCGTAAAAAAGAAGCTGCTGCTAAACTGTCTACCGCTATGTCTGCTGTTG
CTGCTCTGGCTGTTGCTTCTCCGGGTCCGGGTCCGGGTACCGGTACCATCACCGTTGGTG
GTGGTCAGATCAACGGTGTTGGTTTCGCTGCTTACCACTCTACCTACCAGGACAACCTGT
ACGGTCCGGGTCCGGGTTGGGCTGACATCCAGGCTGACCCGGAACGTCGTCGTCGTTACC
AGCAGGGTCCGGGTCCGGGTGCTCCGGAATACGGTGACACCAACGGTTACCGTGCTCCGG
CTGGTCCGGAATACCTGCAGGTTCCGTCTCCGTCTATGGGTCGTGACATCAAAGTTCAGT
TCCAGGAAGCTGTTAACGCTAACGGTATCTACGCTCTGTACCTGCTGACCATGTCTATCG
CTCAGCGTCTGTACGGTTGGGACATCAACGCTACCGAACTGGCTTCTGCTGCTGTTACCG
CTATGGGTTGGTCTCCGTTCTACCCGGCTTTCGAATGGTACGGTGGTCTGGCTGGTTCTA
CCGGTGGTGGTGGTAAAAAAGTTCCGGAATCTGTTGTTATGCCGCCGGGTGAAATCTCTC
TGACCCCGCCGGACGACCTGGGTATCGACATCGCTGCTTACATCACCCCGTCTCGTTGGC
GTATGTACTGGGAAACCTTCCTGACCTCTGAACTGCCGGGTTGGCTGCAGGCTAACCGTC
GTCTGTTCGCTTCTCTGCCGGCTCGTCTGTCTGCTGCTCAGGTTGGTCCGGGTCCGGGTA
AAGTTCCGGAAATCAACGCTGACCGTAAAGAATACCTGCGTTACTCTGCTGTTGTTGGTC
TGTCTATGGCTGCTTCTTCTGCTCTGACCCTGGCTATCTACTCTGACCCGGACGCTGCTC
CGCCGACCTTCTCTGCTCGTCTGACCGCTTACGCTGGTGCTATGCAGCTGTCTATCCCGG
CTATCACCACCCCGGCTTTCACCATCGACCCGGGTCCGACCCTGATCGGTCTGGCTATGG
GTGACGTTATCTCTAACCTGTTCGGTCAGAACGCTCCGGCTATCGCTGCTGCTTCTGACA
TGTGGCGTAAACTGTCTACCGCTATGTCTGCTGTTGCTGCTCTGGCTGTTCCGGCTTGGC
AGCGTAACGACCCGCTGCTGAACGTTGGTAAACTGATCGCTAACATGTCTGCTGTTGCTG
CTCTGGCTGTTGCTGCTTACTACGCTCACACCGACCAGTGGCGTTACCGTGTTTGGGTTT
ACCTGCCGGCTAAATTCCTGGAAGGTTTCGTTCGTACCTCTAACATCAAATTCCAGGACG
CTTACAACGCTGGTGCTGGTGACACCGGTGAACCGACCCCGCAGCGTGCTAACCTGTCTG
ACGCTGCTTACGTTATCGACGCTCTGCCGCTGACCGTTGGTGTTTTCGACTGCGCTGCTA
TCTCTGCTCCGCTGCGTCCGGGTTCTGGTATGCCGCCGTGGGAATACTGGGGTGCTCAGC
TGAACGCTATGAAACCGGACCTGCAGCGTGCTCTGTCTATCGACTTCCGTATGACCTCTA
CCACCCTGGTTTCTGACGTTGGTCCGGGTCCGGGTCGTCGTCGTCGTAAACTGTCTACCG
CTATGTCTGCTGTTGCTGCTGGTCCGGGTCCGGGTATCACCTACATGCGTACCGACTCTA
CCACCCTGTCTGAATCTGCTTAACTCGAG
```
